# Representing the dynamics of natural marmoset vocal behaviors in frontal cortex

**DOI:** 10.1101/2024.03.17.585423

**Authors:** Jingwen Li, Mikio C. Aoi, Cory T. Miller

## Abstract

Here we tested the respective contributions of primate premotor and prefrontal cortex to support vocal behavior. We applied a model-based GLM analysis that better accounts for the inherent variance in natural, continuous behaviors to characterize the activity of neurons throughout frontal cortex as freely-moving marmosets engaged in conversational exchanges. While analyses revealed functional clusters of neural activity related to the different processes involved in the vocal behavior, these clusters did not map to subfields of prefrontal or premotor cortex, as has been observed in more conventional task-based paradigms. Our results suggest a distributed functional organization for the myriad neural mechanisms underlying natural social interactions and has implications for our concepts of the role that frontal cortex plays in governing ethological behaviors in primates.

## Introduction

Natural behaviors typically vary along many dimensions – including their structure, timing, frequency and occurrence with respect to other behaviors. This seemingly indomitable character of ethology has long been a key bottleneck to neuroscience^1^ because of the difficulty it poses to explicating the relationship between the sources of this variance and patterns of neural activity. But as interest in natural behaviors grows, so too does the impetus to overcome these challenges and better understand their neural basis^2–4^. This issue is particularly pertinent to primate frontal cortex – i.e. prefrontal (PFC) and premotor (PMC) - because of the integral role these substrates play in governing a remarkable breadth of complex behavioral and cognitive processes^6,8,10,12,14,16,18–21^. Much of our understanding of primate frontal cortex has been from studies applying conventional, head-fixed task-based paradigms; functions that would reasonably be assumed to be governed by the same subfields when primates engage in analogous natural behaviors. Evidence to this end, however, is not clear. For example, a within-neuron comparison of marmoset PFC activity in both conventional and natural behavior contexts found that responses in the former were not predictive of the latter context, despite the stimuli being identical^5,6^. One potential explanation may be that changes in the state of frontal cortex during natural behaviors^5,7^ or the inherent variability of ethological behaviors^1,4^ may mask the subfield specializations evident in more conventional paradigms. Alternatively, features of natural behaviors – such as the occurrence of varying sequential events^1^ - may drive distinct patterns of neural activity that do not emerge in more focused behavioral tasks^22^. To reconcile these issues, a more refined analysis approach that better accounts for these factors may be needed to characterize patterns of primate frontal cortex activity in natural, ethological behaviors.

Here we applied a Generalized Linear Model (GLM) based approach to characterize neural activity throughout primate frontal cortex – i.e. PFC and PMC – during naturally occurring conversational exchanges in freely-moving marmosets^23,24^ to more accurately characterize the functional organization of these substrates in a natural behavior. Such model-based analyses offer several advantages to this end because they can reveal relationships between neural responses and overlapping external covariates (sensory, motor, state, etc.) along with spike history, and can account for variability in the timing and organization of behaviors thereby capturing facets of brain/behavior interactions that less sophisticated descriptive statistics, such as mean firing rate, cannot^11,13,25,26^. As such, these approaches have the potential to reveal elements of brain function and organization that emerge from dimensions of variability evident in natural behavior that may not be apparent otherwise. Neurophysiological studies of marmoset conversations have shown that populations of PFC and PMC neurons are responsive during different facets of the vocal behavior^15^ including a latent social state that is highly correlated with the probability of conversational exchanges occurring^5,11^. However, the number of neurons whose activity was significantly modulated by these elements of the conversational exchanges were relatively low (∼35%) leaving open the possibility that the more traditional analyses deployed in these studies lacked the statistical power to fully characterize the role that either individual subfields or the population ensembles play in these natural social exchanges. Our analyses demonstrate that the functional organization of this population of cells is modular with respect to function but is spatially distributed.

## Results

### GLMs can characterize continuous events in natural vocal behavior

We recorded the activity of single neurons in PMC and PFC while freely-moving marmosets engaged in their natural vocal behavior (Fig. 1a) and applied different analysis approaches to quantify neural activity, i.e. peri-stimulus time histograms (PSTHs) and GLM based analysis, to determine how these substrates encode the different components of the vocal behavior dynamics. In these experiments, marmosets engaged in conversational exchanges with a computer generated ‘virtual marmoset’ using our interactive playback paradigm, as in previous experiments ^5,27^. To adapt the GLM based analysis to our time-varying, naturalistic experimental paradigm, we included the aforementioned task variables (i.e. behavioral events, internal behavioral states, and spike history) as regressors within a sliding window (Fig. 1b, see Methods & Materials), allowing for a continuous prediction of spike rate over the course of a session (Fig. 1c). The consideration of event history over an entire session has an advantage over simply considering the mean firing rate during specific events as it more effectively considers persistent dynamics over long time periods which typify variable social interactions and can significantly influence neural activity. Behavioral analysis was performed on the continuous recordings to identify vocal-perception events (i.e. hearing calls) and vocal-motor events (i.e. producing calls) as well as the context in which these events occurred (i.e. antiphonal conversation, spontaneous calls, etc.) – and the underlying state (i.e. conversational or spontaneous calling) of the animal^5,17^.

**Figure 1.**
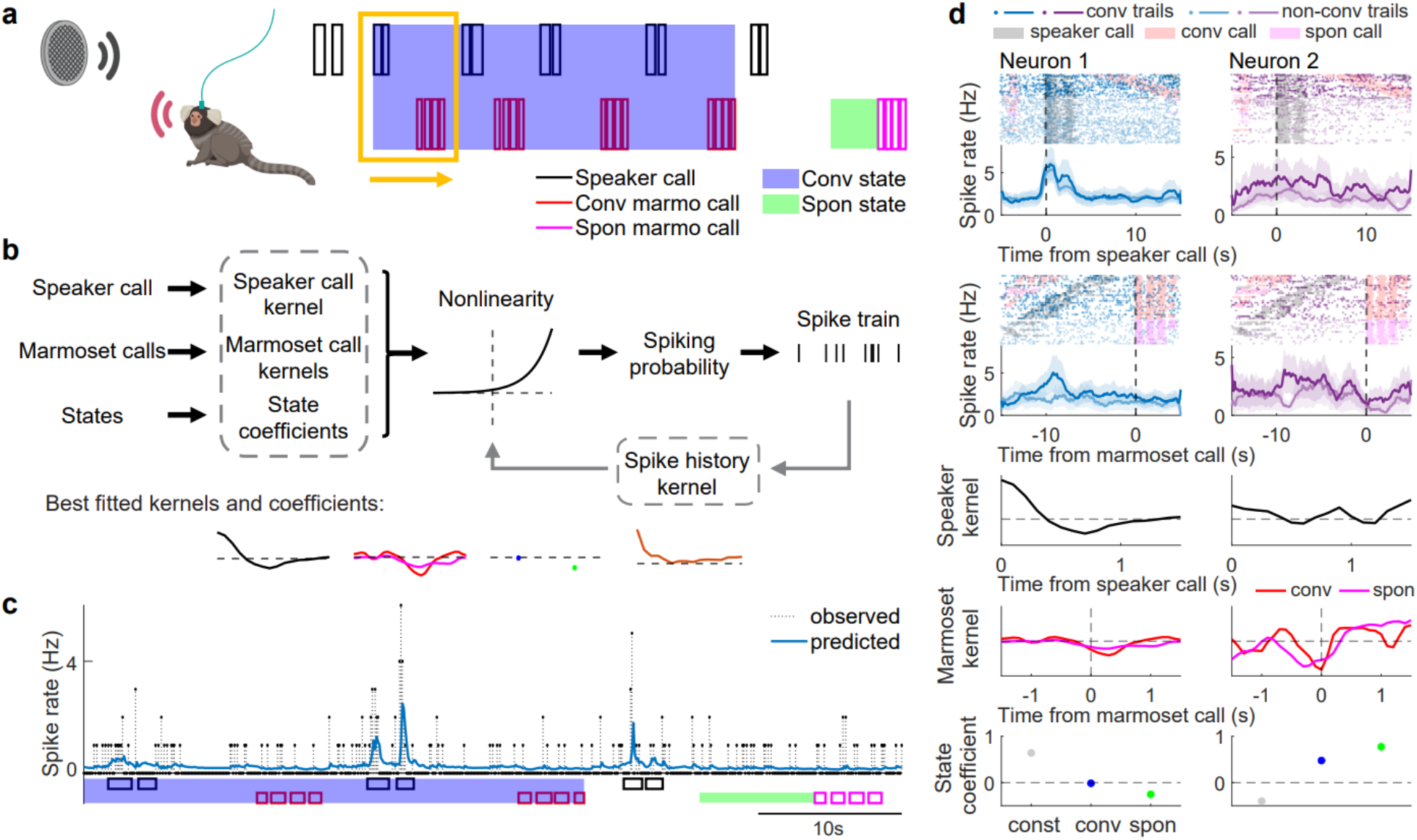
PSTHs and GLM based analysis are applied on activity of neurons in the frontal cortex as marmosets engaged in conversational exchanges. **a)** Experiment setup. Marmosets freely behaved and engaged in natural vocal communication with an interactive playback system. Vocal events and neural activity are simultaneously recorded. Marmoset calls are categorized as conversational (red) and spontaneous calls (pink) depending on whether the marmoset responded to a speak call (black) or initiated a call, respectively. The period during an interactive communication is labeled as conversational state (blue); the period before spontaneous calls is labeled as spontaneous state (green). The yellow box indicates the sliding window used in GLM for the continuous recording. **b)** An illustrative framework of GLM based analysis. Behavioral events, internal states, and spike history are included as regressors in the GLM model to get best fitted kernels and coefficients (in gray dashed box). **c)** Observed and GLM predicted spike rates throughout the recording. The GLM accurately predicts changes in spike rate throughout the recording. **d)** PSTHs and GLM kernels of two example neurons. Neuron 1 exhibited a significant increase in response to hearing a vocalization responded to hearing calls as quantified by the PSTH aligned by conspecific call onset as well as a significant GLM kernel. Neuron 2 exhibited decreased activity during call production in both PSTHs and GLM marmoset call kernels, as well as significant GLM state coefficients quantifying the changes in the activity related to the animal’s conversational and spontaneous states. From top to bottom: spike raster and PSTHs aligned by hearing calls and producing calls, respectively; hearing call kernel; marmoset call kernels (red and pink for conversational and spontaneous call kernels, respectively); state coefficients. In spike raster, darker dots correspond to conversational trails and lighter dots correspond to non-conversational (aligned by hearing calls) or spontaneous (aligned by producing calls) trails; gray, red, pink shaded areas are time period of speaker calls, conversational call, and spontaneous calls, respectively. In PSTHs, darker lines are PSTHs for conversational trails and lighter lines are non-conversational or spontaneous trails; shades represent standard error.

Although previous work has shown that GLM based analyses are effective at modeling spike rates for individual cells in trial-based tasks^7^, we first sought to confirm that our adaptation of this approach to a continuous behavior would yield a similar outcome. As shown in Figure 1d, neuronal firing rates responding to hearing calls or producing calls characterized by GLM kernels were consistent with patterns observed in PSTHs. These results confirmed that our modified GLM analysis could reliably capture multiple facets of continuous, natural primate social interactions (i.e. marmoset conversations).

### GLM encapsulates natural brain/behavior interactions

Building on these observations, we applied both the PSTH and GLM based analysis to each neuron in the population to directly compare how these different approaches encapsulate the various interactions between natural behavior and frontal cortex activity. As highlighted in Figure 2a, we observed that the GLM analysis (81% with permutation test at 0.05 significance level) overwhelmingly outperformed the traditional PSTH analyses (41% with Wilcoxon signed rank test at 0.05 significance level) at identifying neurons with significant modulation by sensory, behavioral, or state events. In fact, the GLM revealed 36% of neurons in the population had significant state-related responses and event-related responses and 19% of neurons had significant state-related responses only, a feature of the natural behavior that was entirely undetected by the PSTH analysis. Analyses revealed that most neurons exhibiting significant changes in activity for hearing and/or producing calls in PSTHs were likewise significant in the GLM analysis, while the opposite was not true (Fig. 2b). We also observed that most of the neurons with significant GLM state coefficients were those with large positive or negative values (Fig. 2c). To more accurately test whether the GLM based analysis was consistent with PSTHs, we used the predicted spike rate from the GLMs to reconstruct PSTHs for both hearing and producing vocalizations (Figure 2d-g). Figure 2d plots the PSTH of a representative neuron when a marmoset heard a conspecific call in both conversational and non-conversational contexts showing that the firing rate predicted by the GLM faithfully recapitulated the PSTH. Though the R^2^ of a neuron was correlated between the conversational and non-conversational contexts, there was no correlation between the R^2^ and the negative log-likelihood of the GLM which represents its overall encoding performance throughout the recording (Figure 2e). This indicates that while the GLM does accurately capture the variability expressed by the PSTHs, it also captures a great deal more. A similar result was observed for vocalization production. As with hearing calls, the call production PSTH was recapitulated by the predicted spike rate from the GLM (Fig. 2f) and the population shows correlated R^2^ of neurons between conversational and spontaneous contexts, while there was no significant correlation between the R^2^ and the negative log-likelihood of the GLM (Fig. 2g). Overall, we observed not only that the predicted spike rate from the GLMs accurately recapitulated the observed PSTHs but that the GLM based analysis was notably more effective at identifying neurons with significant contributions to communication events and state-related processes. As such, the GLM based analysis is a more powerful tool to identify the myriad neural processes that underly dynamic, continuous natural behaviors.

**Figure 2.**
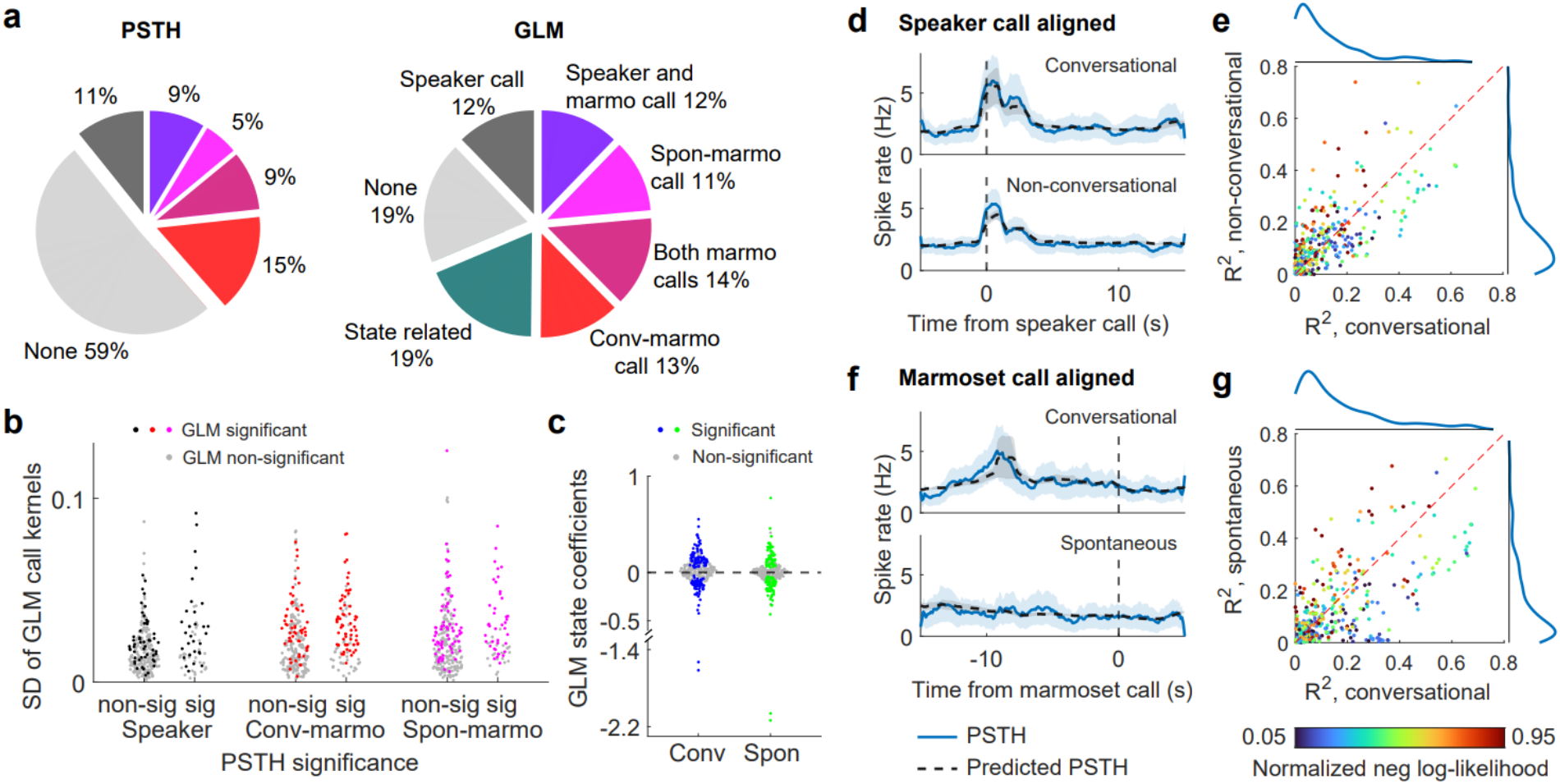
GLM based analysis identifies more neurons with significant functions and successfully recapitulates PSTHs. **a)** Percentage of neurons in categories that significantly respond to communication events or internal behavior states identified by PSTHs and GLMs, respectively. GLM analyses outperformed the traditional PSTH analyses at identifying neurons that significantly respond to behavioral events and internal states. **b)** Standard deviation of GLM kernels for communication events in comparison with PSTH significance. GLMs identified significant neurons responding to behavioral events that are not significant in PSTHs. **c)** GLM coefficients for conversational and spontaneous states. Neurons that significantly respond to internal states are those with large positive or negative values. **d)** Observed PSTH (blue solid line) and predicted PSTH from GLM (black dash line) aligned with hearing calls for an example neuron, grouped by conversational trails (top) and non-conversational trails (bottom). Shaded area represents standard deviation. The predicted PSTHs from GLM well recapitulated the observed PSTHs. **e)** R^2^ between the observed PSTH and the predicted PSTH from GLM of each neuron for non-conversational trails as a function of conversational trails. Blue lines show distribution of R^2^ for the corresponding axes. Color indicates the GLM negative log-likelihood of the neuron normalized to [0,1] among all the neurons. The GLM negative log-likelihood, indicating the overall fitting of GLM throughout the recording, is not correlated to the R^2^ for the hearing event. **f)** Same as in **d** but aligned by producing calls for the same example neuron, grouped by conversational trails (top) and spontaneous trails (bottom). **g)** Same as in **e** but aligned with producing calls.

### Frontal cortex encodes functional clusters

Primate PFC and PMC have long been implicated in a range of higher cognitive functions that, though presumed to be integral to many dynamic natural behaviors^5,7,26,28^, have until recently largely been studied in more traditional, conditioned task paradigms that directly probe specific processes^6,8,10,12,14,16,18^. Applying a model-based analysis to continuous social interactions offers the opportunity to determine the encoding strategy of the primate frontal cortex population for a suite of functions that may covary during natural behavior. Specifically, here we asked whether the marmoset frontal cortex population allocates neurons to distinct functional roles, or do neurons form a continuous spectrum of various functions? To address this question, we collectively analyzed all of the GLM kernels as a single dataset. As a first step we examined the population structure of significant kernels corresponding to the animal hearing a single call using dimensionality reduction (Fig. 3). We applied principal component analysis (PCA) to the GLM kernels and coefficients of all neurons (Fig. 3a). The first 3 PCs explained more than 80% of the variance and the projections of the population on the first two PCs was distributed in PC space with no apparent clustering. Likewise, when PCA was applied to significant marmoset call production kernels the first 4 PCs explained more than 80% of the variance (Fig. 3b). Projections of the population from the first two PCs indicated two clusters, one with both significant conversation and spontaneous call kernels and one with only significant conversation call kernels.

**Figure 3.**
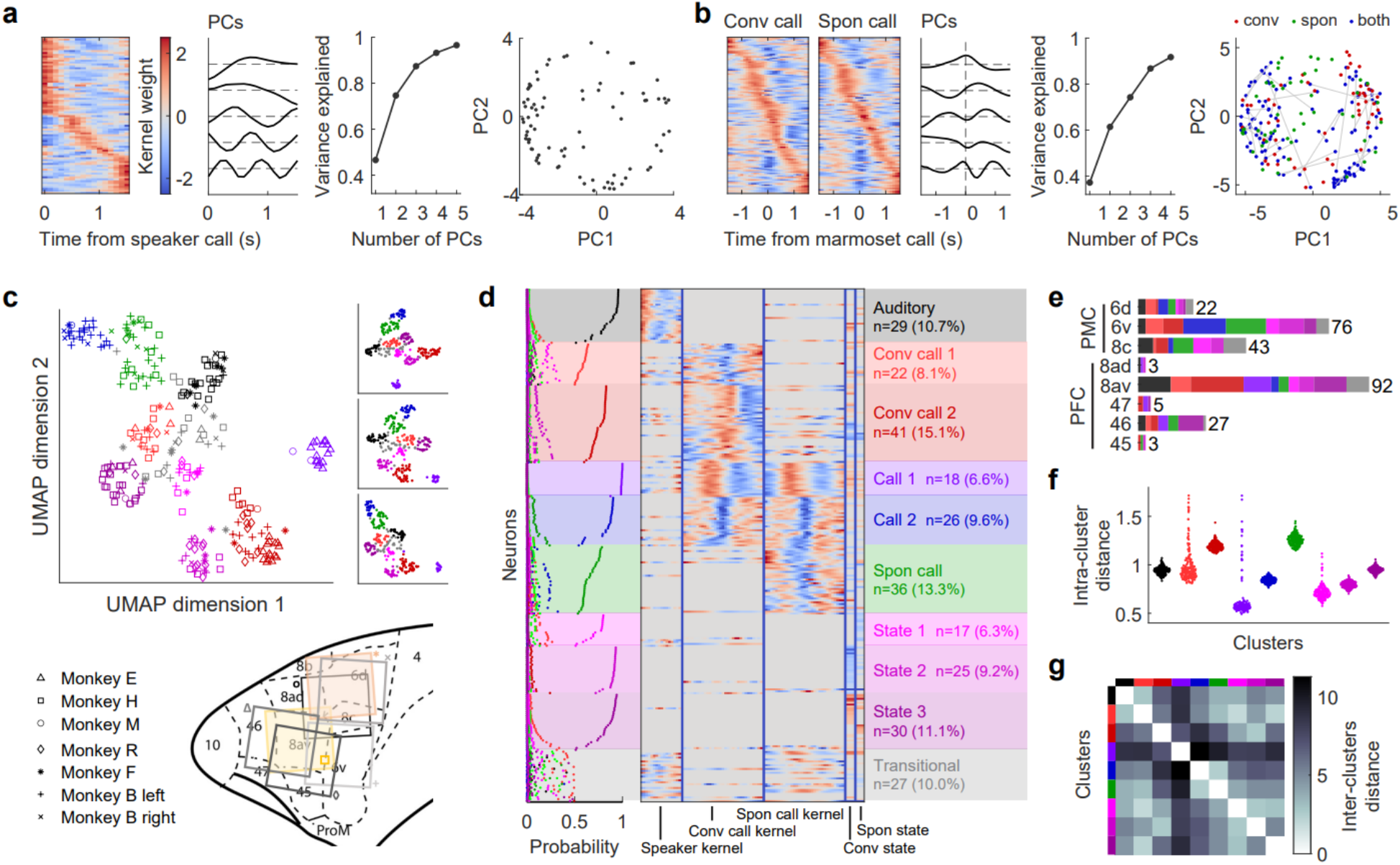
The population in frontal cortex forms a continuous spectrum with clusters of functions from dimensionality reduction and clustering analysis on GLM kernels and coefficients. **a)** PCA of significant speaker kernels. From left to right: significant speaker kernels; corresponding first 5PCs; variance explained by the first 5 PCs; projection of each significant speaker kernel on the first 2PCs. No apparent clustering is observed. **b)** PCA of significant marmoset call kernels. From left to right: significant conversational and spontaneous marmoset call kernels; corresponding first 5PCs; variance explained by the first 5 PCs; projection of each significant kernel on the first 2PCs (units with only significant conversational kernels in red; units with only spontaneous kernels in green; units with both significant conversational and spontaneous kernels in blue and connected by a gray line). Two clusters are observed - one with both significant conversation and spontaneous call kernels and one with only significant conversation call kernels. **c)** Projection of each neuron’s GLM kernels and coefficients on two UMAP dimensions. Each point is a neuron with symbol indicating arrays (position of the array shown below) and color indicating clusters from Gaussian Mixture Model (GMM). The color of the arrays helps to visually distinguish the arrays. The smaller panels show three other examples of UMAP projections. The clustering results were consistent across different UMAP shuffles. **d)** The posterior cluster probability of neurons from GMM (left), the GLM kernels and coefficients of neurons (middle), and the cluster category (right). The clusters are assigned by posterior cluster probability >0.5 and named by their functional roles. Nine clusters were identified with distinguished functional roles and a minority of neurons had ambiguous cluster identity. **e)** Number of neurons recorded in each cortex area where color indicates function. The number on the right indicates the total number of neurons in each area. Neurons in each functional cluster were evident in multiple areas. **f)** Intra-cluster distance for 200 shuffled UMAP projections. The low variance of the intracluster distance indicates the robustness of the clustering results. **g)** Mean inter-cluster distance of 200 shuffled UMAP projections. While one call cluster (purple) shoes high inter-cluster distance with other clusters, the other clusters are grouped with lower inter-cluster distances.

Although subtle clustering features were observed using PCA on individual kernels, we next asked whether the PFC population displayed more distinctive population structure when considering all cognitive functions together. We therefore included kernels corresponding to all communication events and internal states and applied Uniform Manifold Approximation and Projection (UMAP) on the multi-dimensional space formed by the projection of the speaker kernel on the first 3 PCs, the projection of the conversational or spontaneous marmoset call kernels on the first 4 PCs, and the conversational and spontaneous state coefficients (see Methods & Materials). With two UMAP dimensions, we identified distinct clusters using a Gaussian Mixture Model (GMM). The GMM also identified a minority of neurons with ambiguous cluster identity. The clustering results were consistent across different UMAP initializations and final cluster identities were determined by averaging across posterior cluster probabilities from 200 runs of UMAP (Fig. 3c, Methods & Materials). Surprisingly, the functional cluster identity of each neuron was not directly related to where the arrays were positioned in frontal cortex suggesting that functional identities were broadly spatially distributed (Fig. 3c bottom). We identified nine distinct functional clusters: an auditory cluster responding to hearing calls, two clusters responding specifically to producing calls only in conversations, two clusters responding to producing any calls, a cluster responding specifically to producing spontaneous calls, and three clusters mainly associated with the internal states. A minority (10.0%) of neurons had ambiguous cluster identity, with responses that were intermediate between two or more cluster types (Fig. 3d). Neurons in each functional cluster were evident in multiple (on average 5±2) frontal cortex areas (Fig. 3e). For example, neurons in the ‘hearing’ cluster were found in 6 different areas of frontal cortex including premotor and prefrontal areas. To validate the robustness of the clusters, we examined the intra-cluster and inter-cluster distances from the 200 runs of UMAP (Fig. 3f-g). The clusters remained stable across these iterations as indicated by a low variance of intra-cluster distance (Fig. 3f). Inter-cluster distances showed that one call cluster is consistently separable from the other clusters, while other clusters are grouped with lower inter-cluster distances (Fig. 3g).

Overall, these analyses indicate that the primate frontal cortex population has functional clusters which encode the full spectrum of sensory, motor and state related processes that occur during natural vocal communication. Notably, separation of call production into multiple functional clusters - including two each for conversations and two in all contexts – suggests that a more dynamic suite of neural mechanisms underlies this facet of vocal behavior that is not apparent in our broad behavioral classifications. A more refined consideration of the ethology itself is likely needed to better characterize the suite of computational processes underlying the behavior.

### Natural behavior can be predicted from GLM kernels

The distinct functional clusters encoded by marmoset frontal cortex are suggestive of the broad role that these neocortical substrates play in supporting natural social interactions. To further explore the significance of these clusters, we next tested the respective contributions of neurons in these functional clusters to predict whether a marmoset will produce a vocalization in response to hearing a conspecific call. Because kernels in each cluster varied in their sensory and vocal-motor related activity, as well as context and state, it is possible that the activity of only some neurons in particular clusters carry predictive information about the subjects’ vocal behavior. Each time a conspecific call was heard, we calculated a conversational call score (CCS) corresponding to the log-likelihood-ratio of a GLM that assumed a marmoset emitted a conversational call – defined as a call produced in response to hearing a conspecific call – versus one assuming no response (Fig. 4a; see Method & Materials). If the marmoset produced a conversational call after hearing a call, the GLM assuming a call occurred should have a better fit to the data and yield a high CCS. If the marmoset did not respond, the GLMs should have a worse fit when assuming a conversational call exists and yield a low CCS. We selected the highest score after the marmoset hears a call to represent how likely the marmoset was to engage in the conversation and calculated the Receiver Operating Characteristic (ROC) curve and Area Under Curve (AUC) for each neuron (Fig. 4a). This analysis revealed a range of decoding performances among marmoset frontal cortex neurons, where 32.5% of neurons yielded significant decoding performance with some neuron’s AUC as high as 0.8 and above (Fig. 4b) indicating robust predictability of this behavioral event. The decoding performance of neurons in each functional cluster is not significantly different from each other, and neurons with high decoding performance were distributed across clusters (Fig. 4c). Likewise, the population of neurons with the highest decoding performance consisted of neurons with various functions (Fig. 4d). Together these results further support our observation that distinct regions of marmoset frontal cortex do not exhibit distinct functions, but rather that the ensemble population activity contributes to different facets of natural vocal communication.

**Fig 4.**
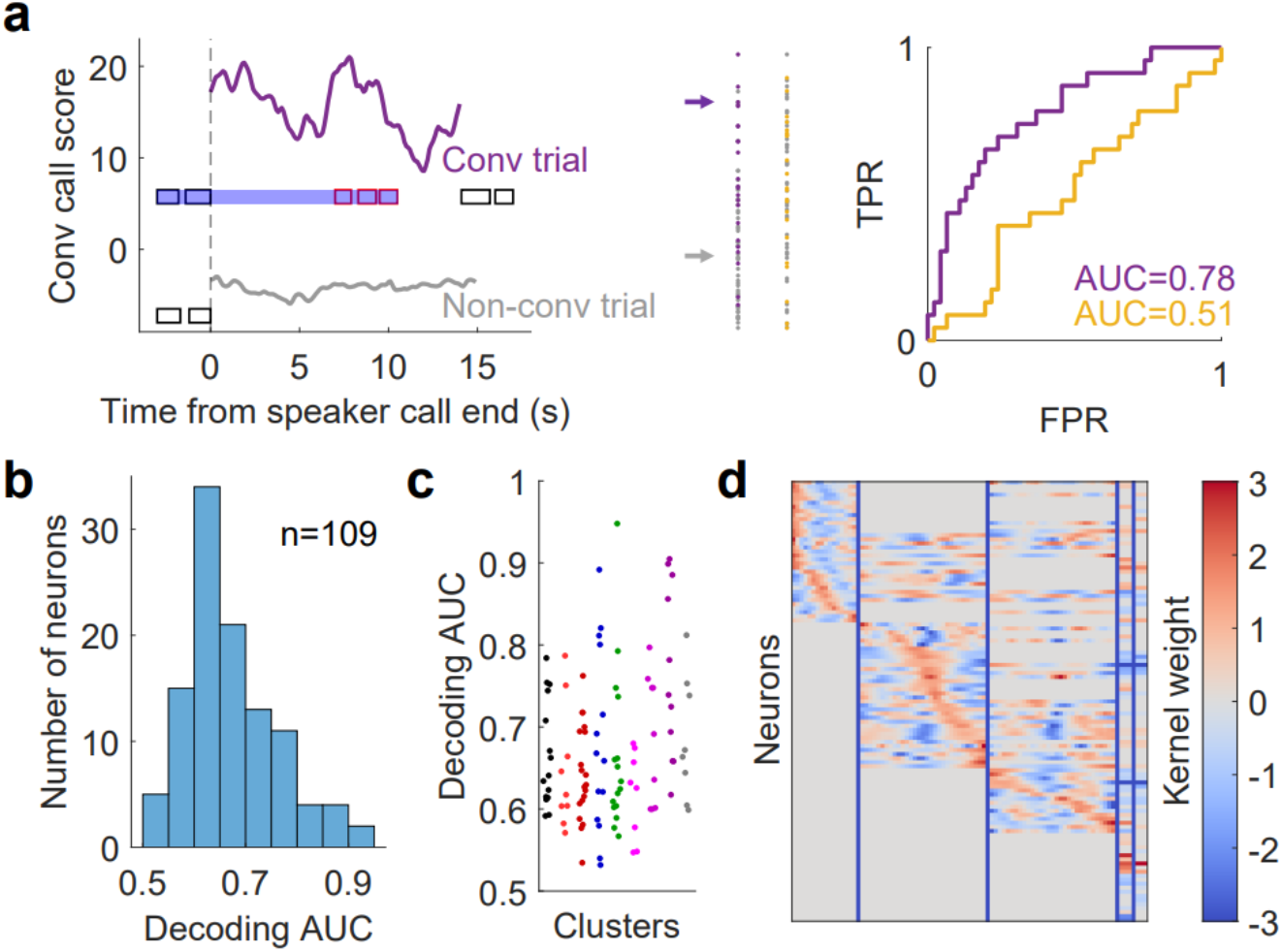
Using GLMs to decode a marmoset’s response is successful for neurons from various clusters and functions. **a)** Decoding process illustrated by two example neurons. Left: conversational call score measured by GLM negative log-likelihood as a function of time from speaker call end for a conversational trial (purple line) and a non-conversational trial (gray line) of a neuron. Middle: conversational call scores (CCS) for all trials of two neurons (purple and yellow). Each dot represents a score of a trial. Dots in purple or yellow are conversational trials and gray trials. The two arrows represent the two trials in the left panel. Right: ROC curve and AUC of the two neurons in the middle panel. **b)** Distribution of decoding AUC for neurons with significant GLM kernels or coefficients and significant decoding performance. 109 neurons yielded significant decoding performance. **c)** Decoding AUC of neurons grouped in different clusters. Color of the clusters is the same as in Fig. 3. Neurons with significant decoding performance are distributed in various clusters. **d)** GLM kernels and coefficients of the neurons with significant decoding performance. The population with high decoding performance consists of neurons with various functions.

## Discussion

Here we examined the respective contributions of primate PFC and PMC to processes underlying marmoset conversations to explicate how single neurons and population ensembles govern this continuous natural behavior. Given the variability inherent to the myriad behavioral processes that occur during marmoset conversational exchanges, we adapted a GLM based analysis^19,20^ to evaluate the activity of neurons throughout frontal cortex as marmosets engaged in their natural vocal interactions. Our model-based GLM analysis robustly outperformed more traditional PSTH based analyses, as it identified more neurons with significant vocal behavior related functions – i.e. hearing and producing calls in different social contexts - as well as state-related neural activity. The GLM analysis also revealed novel functional clusters in marmoset frontal cortex encapsulating the different behavioral and state related properties of the vocal behavior.

Importantly, these functional clusters were distributed in an anatomically heterogenous organization that had not previously been observed in more traditional head-fixed task-based experiments, including those testing vocalization recognition and production^23,29–32^. These results suggest that primate frontal cortex activity is involved in nearly all facets of natural, continuous vocal behaviors through an anatomically distributed - but functionally modular - pattern of ensemble activity.

Our analysis approach innovated on previous GLM-based applications to neurophysiological data analysis in two important ways that may be integral to generalizing this approach to other natural behaviors. First, we used a moving-window to study the encoding properties of cells on a moment-by-moment, but continuous basis. Second, rather than only inferring and interpreting the learned kernels for each cell independently we also took a population-level view of the cell encoding properties. By performing dimensionality reduction and clustering analysis on the GLM kernels and coefficients, we revealed clusters in the population playing different functional roles that spanned the breadth of the natural behavior as an ensemble code across the population. Notably, these clusters were not localized to different sub-fields of frontal cortex, but rather formed a continuum of functional clusters that were largely independent of their respective anatomical position. This population encoding strategy may benefit the system in maximizing information propagation or functional robustness as has been recently shown in primary visual cortex^29^ and retrosplenial cortex^33^. A balance of segregation and integration must be maintained in frontal cortex to flexibly adapt cognitive strategies, a particularly powerful computational mechanism for mediating social interactions that vary due to - sometimes unpredictable - changes in the social and ecological landscapes. While this pattern is different from previous observations of mixed-selectivity in marmoset PFC across different contexts^15^, neural activity in the current study was only recorded during natural communication. It’s possible that the functional role of these neurons may be consistent within a particular context but change as a function of the demands of the immediate environment.

Perspectives on the neural basis of primate behavior are evolving in hand with the increased use of ethological paradigms^22,26,28,34,35^ and particularly as effects in conventional tasks are not necessarily recapitulated in the presumptive analogous natural behaviors^15,32^. Our results here showed that - while functional clusters of these processes are evident in the prefrontal and premotor cortices of marmosets engaged in natural vocal interactions - the organization is widely distributed throughout frontal cortex; a stark contrast to the most directly analogous studies with conventional paradigms^29,30,32^. We conjecture that the distributed functional organization during natural marmoset conversations was evident because the behavioral and cognitive processes that have traditionally been studied selectively through more targeted experimentation become more covaried in continuous, natural behaviors and, as such, rely on a more distributed computational ensemble coding strategy in primate frontal cortex^22^. Though admittedly limited in number, the few studies to directly compare neural responses in the primate brain between conventional and naturalistic paradigms have reported considerable differences^15,32^ that may portend the need to evolve our conceptions of brain computations and behavior. Rather than be an impediment to understanding frontal cortex function - or the brain more generally - the behavioral variability and covariance inherent to ethological behaviors may be a uniquely powerful engine of discovery, as it is only in these contexts that at least some computational mechanisms needed to govern the more distinct primate cognitive faculties may emerge and be open to scientific inquiry^36–38^.

## Acknowledgements

We thank Drs. Vlad Jovanovic, Sam Nummela and Wren Thomas for assistance with data collection. This work was supported by NIH R01 DC012087.

## STAR*METHODS

## KEY RESOURCES TABLE

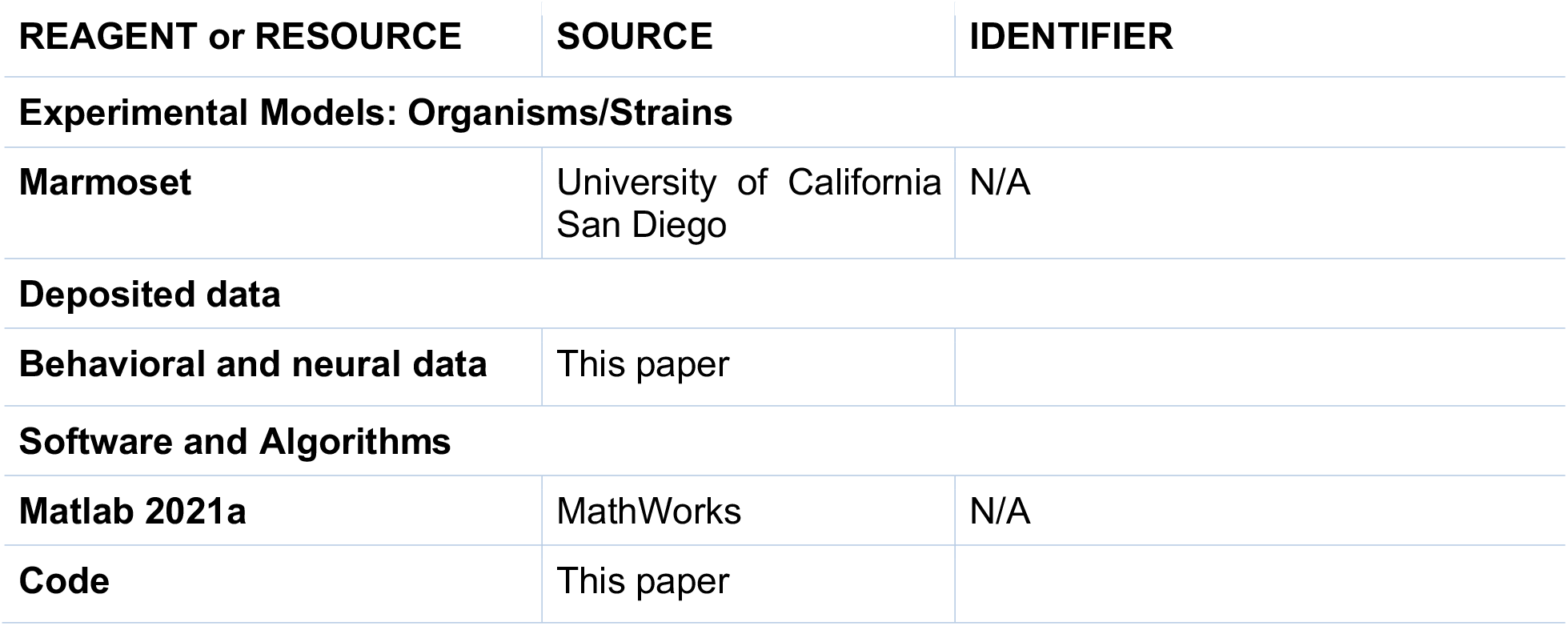

## RESOURCE AVAILABILITY

### Lead contact

Further information and requests for resources should be directed to and will be fulfilled by the lead contact, Cory Miller.

## Materials availability

This study did not generate new unique reagents.

## Data and code availability

Data and code are available to the public (see key resources table).

## EXPERIMENTAL MODEL AND SUBJECT DETAILS

### Subjects

Experiments in this work were done with six adult common marmosets (*Callithrix jacchus*). Monkey E, F, H and M were female and monkey B and R were male. All subjects were group housed and were at least 1.5 years old at time of implant. Monkey B had bilateral arrays implanted and all the other monkeys had a single array implanted. All experiments were performed in the Cortical Systems and Behavior Laboratory at University of California San Diego (UCSD) and approved by the UCSD Institutional Animal Care and Use Committee.

### Experiment model

The recordings took place in a 4×3 m test room. Subjects were placed in a 32x18x46 cm test cage that allows them to freely behave and engage in natural vocal communication behaviors, as in previous behavioral and neurophysiological experiments^5,27,39^. We employed an interactive playback software that responses to subjects’ calls and simulates natural vocal interactions. When subjects produce a phee call, a responsive phee call was broadcasted in response to the subject’s phee call within 2-4 s after the subjects’ phee call offset. If subjects produce no phee calls for 45-60 s, a phee call was broadcasted to initiate a vocal interaction. Phee calls from subjects within 10 s after a broadcasted phee call are classified as a conversational response; subjects’ phee calls outside the time window are classified as spontaneous phee calls. Detailed description of the behavioral paradigm in this work can also be found in the previous work Jovanovic et al. 2019 for subjects H, E and M, and Miller et al. 2015 for subjects F, B and R.

## METHOD DETAILS

### Neurophysiology

The surgical procedure of the chronic microelectrode array implantation is described in our previous work (subjects H, E and M ^15^;and subjects F, B and R ^23^. The microelectrode array consists of 16 channel tungsten electrodes each housed in an independent guide tube in a 4*4 mm grid (Neuralynx, Bozeman, MT). Each array was positioned on the surface of the brain in the chronic implant, and perpendicularly lowered to the laminar surface of neocortex with a calibrated Warp Drive pusher attached to the end of the guide tubes. The electrode was advanced 10-20 um twice a week over the course of the experiment sessions. The headstage preamplifier was connected to the arrays and attached to a tether to allow subjects to freely move in the experiment box. The tether was wrapped by a metal coil to prevent interference from the movement and vocalization of the subjects. Spike sorting was done off-line with 300-9000 Hz filtering, thresholding, 1 ms waveform PCA, DBSCAN automatic clustering and manual curation. Single units were determined with the criteria that the signal-to-noise ratio >= 13 dB and less than 1% inter-spike intervals violate the refractory period < 1 ms. Overall, 335 single units met the criteria and were studied in this work.

## QUANTIFICATION AND STATISTICAL ANALYSIS

### PSTH analysis

PSTHs were plotted using 0.1 s time bins with a boxcar average filter on 15 data points. The significance of neural responses in PSTH was tested on the firing rate during 2 s peri-stimulus time versus firing rate during stimulus with the Wilcoxon signed rank test at 0.05 significance level.

### GLM based analysis

The GLM based analysis is built upon the model:

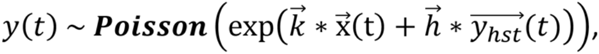

where *y(t)* is spike count at time *t, y*_hst_ is the history of spiking up to time *t, x* represents behavioral events and internal behavioral states, *h* and *k* represent the kernels or coefficients for the events or states, respectively. Behavioral events include hearing a call, producing a call in response to hearing a conspecific call, and producing a spontaneous call; internal behavioral states include conversational state, defined as the period that the marmoset is engaged in vocal interactions, and spontaneous state, defined as 8 s before the marmoset emits a spontaneous call. The spike trains and behavioral data were discretized into 0.01 s bins. Thus, spike trains were converted to spike count time series, and behavior events and states were converted to binary time series as input to the model. A sliding window was used to collect samples at different time *t*. We used the canonical exponential nonlinearity for the activation function *f* and obtained kernels and coefficients by maximizing the log posterior likelihood with a smoothing prior:

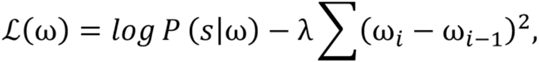

where *s* is the spike rate in data, *w* represents the kernel coefficients, and *λ* is the penalty parameter. Notably, the regularization term penalizes differences in neighboring kernel coefficients and encourages smoothness of the kernel. This formulation is equivalent to a choosing a prior over the differences in neighboring coefficients as independent Gaussian random variables with zero mean and variance 1/*λ*. For each neuron, *λ* was determined by fivefold cross-validation (Fig. S1). Significance of the kernels and coefficients were determined by permutation test at 0.05 significance level. The null kernels and coefficients in the permutation test for each neuron were obtained by shuffling the alignment of spike trains and behavioral data 1000 times. This null model breaks the temporal association between spikes and behavior without altering the temporal autocorrelations of the time series. For the kernels, the permutation test compares the peak of each kernel with its corresponding null distribution; for the coefficients, the permutation test is performed on the coefficient and null distribution of coefficients.

### Clustering analysis

Clustering analysis of the population was applied on the neurons with at least 1 significant kernel or state coefficient. For each neuron, we took the projections on the first 3 PCs of the speaker call kernel, the first 4 PCs of the two marmoset call kernels, the conversational and the spontaneous coefficients to form a 13-dimensional space. The number of PCs used is determined by 80% variance explained by the PCs. We found that higher numbers of included PCs obscured the shared features among kernels and diminished clustering performance, while fewer PCs omitted important features to capture clustering structure. UMAP was applied to the PC loadings for nonlinear dimensionality reduction.

A Gaussian Mixture Model (GMM) with 10 components was applied on the two UMAP dimensions. For each neuron and run of UMAP, fitting the GMM gives a posterior probability of cluster memberships. To account for variability in cluster geometry across runs of UMAP, we matched cluster identity across runs of UMAP by minimizing the Wasserstein-2 distance between clusters and then averaged posterior cluster membership across all 200 UMAPs. This procedure is equivalent to approximate marginalization across the ensemble of UMAP runs. We assigned a neuron to a cluster if there exists one dominant component (average posterior probability > 0.5).

Otherwise, the neuron is considered as transitional which lies in between clusters. To test the robustness of the clustering analysis, intra-cluster and inter-cluster distances are defined as the average distance of the nodes within the cluster and the distance between the center of two clusters, respectively.

### Decoding

We defined a conversational call score (CCS) to represent how likely a call is produced by the marmoset after hearing a call. At each instance of a conspecific call was heard, we fit 2 models; one where we assumed that the marmoset produced a call with 4 s duration at time *t* after the offset of the conspecific call was heard, and one where no call was produced. Each version of the model was fit with a GLM which yielded 2 series of log-likelihood values for *t* from 0-10 s or until another conspecific call was broadcasted to the marmoset with 0.1 s time bin. The CCS is defined as the maximum log-likelihood-ratio of the two models at the time after the conspecific call offset. Ground truth of the marmoset vocal behavior is used to calculate ROC and AUC of each neuron’s decoding performance. Significance of the decoding performance are determined by permutation test at 0.05 significance level. The null decoding performance in the permutation test is obtained by shuffling the alignment of spike trains and behavioral data 1000 times.

**Figure S1.**
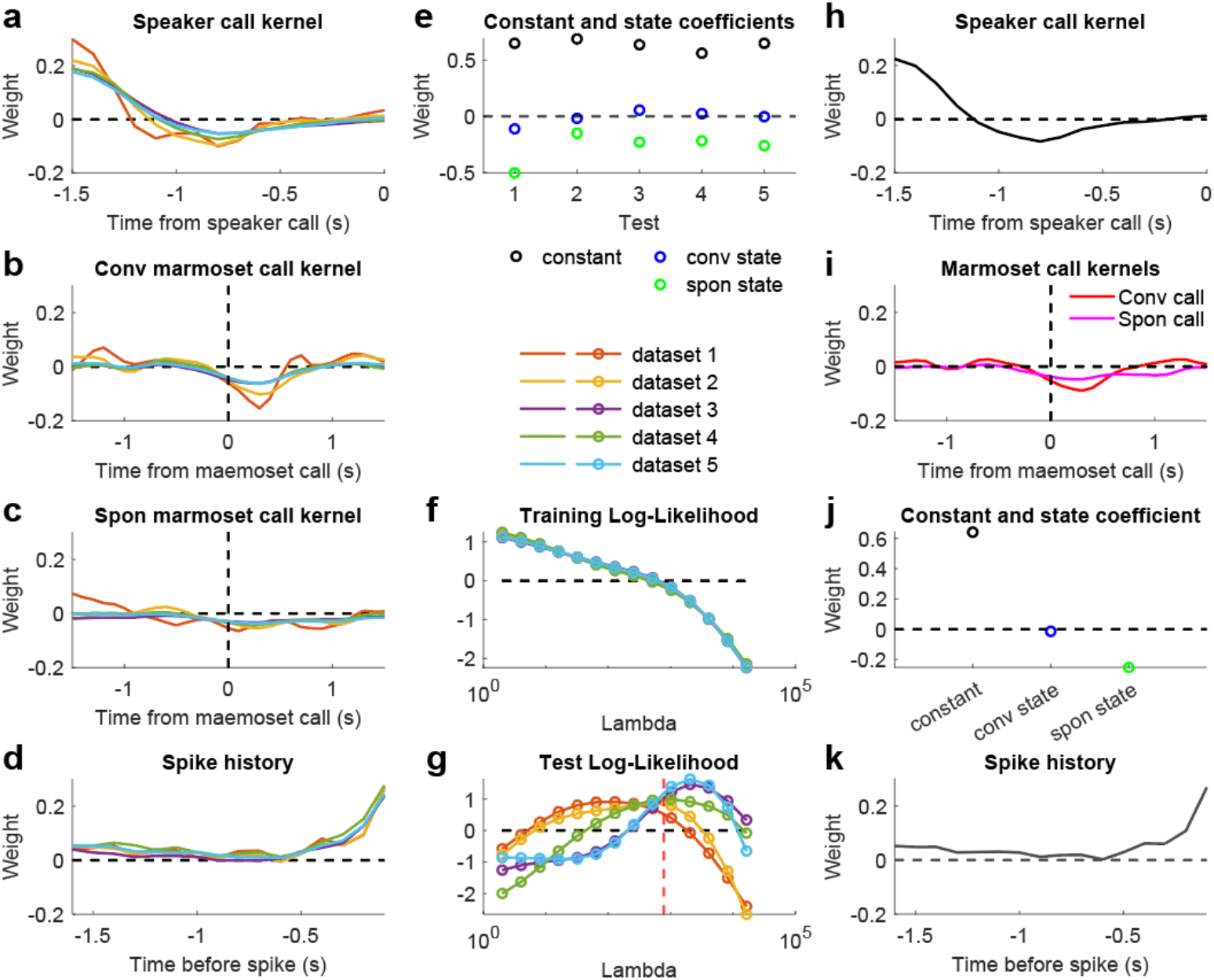
Cross-validation in GLM for an example neuron. **a-e)** Kernels and coefficients for fivefold cross-validation. **f)** Log-likelihood obtained from GLM in each training set as a function of the penalty parameter lambda. The Log-likelihood drops as lambda increases. **g)** Log-likelihood obtained from GLM in each test set as a function of the penalty parameter lambda. The Log-likelihood reaches an optimal value at different lambda for different test set. The optimal lambda (red dash line) is chosen at the logarithmic average of the five datasets. **h-k)** Kernels and coefficients obtained with the optimal lambda for the whole dataset.

## Literature Cited

1. Miller, C.T., Gire, D., Hoke, K., Huk, A.C., Kelley, D., Leopold, D.A., Smear, M.C., Theunissen, F., Yartsev, M., and Niell, C.M. (2022). Natural behavior is the language of the brain. Curr. Biol. 32, R482–R493.

2. Datta, S.R., Anderson, D.J., Branson, K., Perona, P., and Leifer, A. (2019). Computational Neuroethology: A Call to Action. Neuron 104, 11–24.

3. Dennis, E.J., El Hady, A., Michaiel, A., Clemens, A., Tervo, D.R.G., Voigts, J., and Datta, S.R. (2021). Systems Neuroscience of Natural Behaviors in Rodents. J. Neurosci. 41, 911–919.

4. Pereira, T.D., Shaevitz, J.W., and Murthy, M. (2020). Quantifying behavior to understand the brain. Nat. Neurosci. 23, 1537–1549.

5. Jovanovic, V., Fishbein, A.R., de la Mothe, L., Lee, K.-F., and Miller, C.T. (2022). Behavioral context affects social signal representations within single primate prefrontal cortex neurons. Neuron. 10.1016/j.neuron.2022.01.020.

6. Miller, E.K., Lundqvist, M., and Bastos, A.M. (2018). Working memory 2.0. Neuron 100, 463–475.

7. Nummela, S., Jovanovic, V., de la Mothe, L.A., and Miller, C.T. (2017). Social context-dependent activity in marmoset frontal cortex populations during natural conversations. Journal of Neuroscience 37, 7036–7047.

8. Lara, A.H., and Wallis, J.D. (2015). The Role of Prefrontal Cortex in Working Memory: A Mini Review. Front. Syst. Neurosci. 9, 173.

9. Miller, C.T., Thomas, A.W., Nummela, S.U., and de la Mothe, L.A. (2015). Responses of primate frontal cortex neurons during natural vocal communication. J. Neurophysiol. 114, 1158–1171.

10. Miller, E.K., Nieder, A., Freedman, D.J., and Wallis, J.D. (2003). Neural correlates of categories and concepts. Curr. Opin. Neurobiol. 13, 198–203.

11. Park, I.M., Meister, M.L.R., Huk, A.C., and Pillow, J.W. (2014). Encoding and decoding in parietal cortex during sensorimotor decision-making. Nat. Neurosci. 17, 1395–1403.

12. Mansouri, F.A., Freedman, D.J., and Buckley, M.J. (2020). Emergence of abstract rules in the primate brain. Nat. Rev. Neurosci. 21, 595–610.

13. Latimer, K.W., Yates, J.L., Meister, M.L.R., Huk, A.C., and Pillow, J.W. (2015). Single-trial spike trains in parietal cortex reveal discrete steps during decision-making. Science. 10.1126/science.aaa4056.

14. Hernandez, A., Zainos, A., and Romo, R. (2002). Temporal evolution of a decision-making process in medial premotor cortex. Neuron 33, 959–972.

15. Parker, P.R.L., Abe, E.T.T., Leonard, E.S.P., Martins, D.M., and Niell, C.M. (2022). Joint coding of visual input and eye/head position in V1 of freely moving mice. Neuron 110, 3897-3906.e5.

16. Wallis, J.D., and Miller, E.K. (2003). From rule to response: neuronal processes in the premotor and prefrontal cortex. J. Neurophysiol. 90, 1790–1806.

17. McMahon, D.B., Russ, B.E., Elnaiem, H.D., Kurnikova, A.I., and Leopold, D.A. (2015). Single-Unit Activity during Natural Vision: Diversity, Consistency and Spatial Sensitivity among AF Face Patch Neurons. Journal of Neuroscience 35, 5537–5548.

18. Buschman, T.J. (2021). Balancing flexibility and interference in working memory. Annu. Rev. Vis. Sci. 7, 367–388.

19. Fuster, J.M. (2008). The Prefrontal Cortex (Academic Press).

20. Petrides, M. (2005). Lateral prefrontal cortex: architectonic and functional organization. Philos. Trans. R. Soc. Lond. B Biol. Sci. 360, 781–795.

21. Wise, S. (1985). The primate premotor cortex: Past, present, and preparatory. Annu. Rev. Neurosci. 8, 1–19.

22. Testard, C., Tremblay, S., Parodi, F., DiTullio, R.W., Acevedo-Ithier, A., Gardiner, K.L., Kording, K., and Platt, M.L. (2024). Neural signatures of natural behaviour in socializing macaques. Nature. 10.1038/s41586-024-07178-6.

23. Grijseels, D.M., Prendergast, B.J., Gorman, J.C., and Miller, C.T. (2023). The neurobiology of vocal communication in marmosets. Ann. N. Y. Acad. Sci. 1528, 13–28.

24. Grijseels, D.M., Fairbank, D.A., and Miller, C. (2023). A model of marmoset monkey vocal turn-taking. 10.2139/ssrn.4600721.

25. Aoi, M.C., Mante, V., and Pillow, J.W. (2020). Prefrontal cortex exhibits multidimensional dynamic encoding during decision-making. Nat. Neurosci. 23, 1410–1420.

26. Voloh, B., Maisson, D.J.-N., Cervera, R.L., Conover, I., Zambre, M., Hayden, B., and Zimmermann, J. (2023). Hierarchical action encoding in prefrontal cortex of freely moving macaques. Cell Rep. 42, 113091.

27. Miller, C.T., and Wren Thomas, A. (2012). Individual recognition during bouts of antiphonal calling in common marmosets. J. Comp. Physiol. A Neuroethol. Sens. Neural Behav. Physiol. 198, 337–346.

28. Maisson, D.J.-N., Cervera, R.L., Voloh, B., Conover, I., Zambre, M., Zimmermann, J., and Hayden, B.Y. (2023). Widespread coding of navigational variables in prefrontal cortex. Curr. Biol. 33, 3478-3488.e3.

29. Romanski, L.M., Averbeck, B.B., and Diltz, M. (2005). Neural representation of vocalizations in the primate ventrolateral prefrontal cortex. J. Neurophysiol. 93, 734–747.

30. Hage, S.R., and Nieder, A. (2013). Single neurons in monkey prefrontal cortex encode volitional initiation of vocalizations. Nat. Commun. 4, 2409.

31. Jafari, A., Dureux, A., Zanini, A., Menon, R.S., Gilbert, K.M., and Everling, S. (2023). A vocalization-processing network in marmosets. Cell Rep. 42, 112526.

32. Plakke, B., Ng, C.W., and Poremba, A. (2013). Neural correlates of auditory recognition memory in primate lateral prefrontal cortex. Neuroscience 244, 62–76.

33. Sit, K.K., and Goard, M.J. (2023). Coregistration of heading to visual cues in retrosplenial cortex. Nat. Commun. 14, 1992.

34. Shaw, L., Wang, K.H., and Mitchell, J. (2023). Fast prediction in marmoset reach-to-grasp movements for dynamic prey. Curr. Biol. 33, 2557-2565.e4.

35. Mao, D., Avila, E., Caziot, B., Laurens, J., Dickman, J.D., and Angelaki, D.E. (2021). Spatial modulation of hippocampal activity in freely moving macaques. Neuron 109, 3521-3534.e6.

36. Miller, E.K., and Cohen, J.D. (2001). An integrative theory of prefrontal cortex function. Annu. Rev. Neurosci. 24, 167–202.

37. Martinez-Trujillo, J. (2022). Visual attention in the prefrontal cortex. Annu. Rev. Vis. Sci. 10.1146/annurev-vision-100720-031711.

38. Miller, E.K. (2000). The prefrontal cortex and cognitive control. Nat. Rev. Neurosci. 1, 59–65.

39. Miller, C.T., Iguina, C.G., and Hauser, M.D. (2005). Processing vocal signals for recognition during antiphonal calling in tamarins. Anim. Behav. 69, 1387–1398.

